# A novel intranasal administration adenoviral vector-based platform for rapid COVID-19 vaccine development

**DOI:** 10.1101/2022.02.21.481247

**Authors:** Yifei Yuan, Xing Gao, Fengfeng Ni, Wenbo Xie, Wenbin Fu, Gaoxia Zhang, Huimin Hu, Yuncheng Li, Qinxue Hu, Chuanmengyuan Huang, Bo Liu, Yalan Liu, Qiong Shen, Min Liang

**Author notes:** Corresponding authors. *E-mail addresses* (Y. Liu), (Q. Shen) or (M. Liang). Yifei Yuan, Xing Gao and Fengfeng Ni have contributed equally to this work.

## Abstract

The coronavirus SARS-CoV-2 has a severe impact on global public health, and the emerging variants threaten the efficacy of the circulating vaccines. Here, we report that a single vaccination with a non-replicating chimpanzee adenovirus-based vaccine against the SARS-CoV-2 Delta variant (JS1-delta) elicits potent humoral, cellular and mucosal immunity in mice. Additionally, a single intranasal administration of JS1-delta provides effective protection against the Delta (B.1.617.2) variant challenge in mice. This study indicates that chimpanzee adenovirus type 3 (ChAd3) derived vector represents a promising platform for antiviral vaccine development against respiratory infections and JS1-delta is worth further investigation in human clinical trials.

**Highlights:**

1. A new chimpanzee adenoviral vaccine against the SARS-CoV-2 Delta variant was developed.
2. The vaccine elicited potent humoral, cellular and mucosal immunity in mice.
3. The vaccine protected mice from the Delta variant challenge.

## 1. Introduction

The global coronavirus disease 2019 (COVID-19) pandemic caused by the severe acute respiratory syndrome coronavirus 2 (SARS-CoV-2) has profound impacts on public health and economic life. As of 23 June 2022, there have been over 539 million laboratory-confirmed COVID-19 cases and more than 6.3 million conformed deaths (WHO, 2022a). In order to prevent this disease, there are at least 21 COVID-19 vaccines from different platforms have been approved worldwide for emergency use (Rahman et al., 2022). Among them, nine vaccines are recommended by the World Health Organization (WHO) (WHO, 2022d). These nine vaccines can be divided into four types, including the inactivated whole-virion (BBIBP-CorV, CoronaVac and COVAXIN), non-replicating viral vector (ChAdOx1 nCoV-19, Ad26.COV2.S and Ad5-nCoV), protein subunit (NVX-CoV2373), and nucleic acid (BNT162b2 and mRNA-1273) vaccines (Rahman et al., 2022; WHO, 2022d). Although the currently used vaccines have been demonstrated to reduce severe disease and death effectively, the constantly emerging SARS-CoV-2 variants, such as Delta (B.1.617.2) and Omicron (B.1.1.529) which have spread quickly around the world, pose serious threats to existing immunologic barriers by compromising the neutralization capability of antibodies and causing breakthrough infection (Hacisuleyman et al., 2021; Ma et al., 2021; Tian et al., 2021). Therefore, new vaccines against the pandemic variants should be urgently needed.

Recombinant adenoviral vectors (rAds) are powerful inducers of cellular immune responses and have been highly attractive for vaccine development and gene therapy (Geisbert et al., 2011). Adenoviruses (Ads) are about 65-90 nm in size, and contain a 26-48 kb non-segmented linear double-stranded DNA genome with the inverted terminal repeats (ITRs) located on both ends (Kulanayake and Tikoo, 2021; Russell, 2009; Saha et al., 2014). According to the time of expression, the adenoviral coding regions can be classified into early regions (E1-E4) and late regions (L1-L5) (Davison et al., 2003; Saha et al., 2014). The E1 region initiates the viral life cycle and encodes proteins necessary for the expression of other viral genes (Kovesdi and Hedley, 2010; Saha et al., 2014). The E3 region encodes proteins that counteract the host immune responses, but is not essential for viral replication *in vitro* (Horwitz, 2004; Kovesdi and Hedley, 2010; Wold et al., 1995). Therefore, the first generation of rAds is commonly engineered by replacing the E1 region with transgene cassettes and removing the E3 region to free up more space (Bulcha et al., 2021; Kovesdi and Hedley, 2010). The E1/E3 deleted rAds are replication-incompetent, but can be produced to high titers in producer cells, such as HEK293 and PER.C6 cell lines (Geisbert et al., 2011; Kovesdi and Hedley, 2010). With an insertion of the human adenovirus type 5 (hAd5) E1 region into their chromosomes, the producer cells can support the replication of the first-generation adenoviral vectors (Fallaux et al., 1998; Kovesdi and Hedley, 2010; Louis et al., 1997). Although new generations of rAds are under continuous development, the first-generation is still widely used in preclinical and clinical researches (Cots et al., 2013; Palmer and Ng, 2005; Wang and Finer, 1996). There are several types of adenoviral vector COVID-19 vaccines have been approved for emergence use, including hAd5 (the second component of Sputnik V developed by the N.F. Gamaleya Center and Ad5-nCoV developed by CanSino), hAd26 (Ad26.COV2.S developed by Janssen and Sputnik Light developed by the N.F. Gamaleya Center) and ChAdY25 (ChAdOx1 nCoV-19 developed by Oxford/AstraZeneca) (Bos et al., 2020; Logunov et al., 2020; Rahman et al., 2022; van Doremalen et al., 2020; Zhu et al., 2020). As the progress of the COVID-19 pandemic, more and more new types of adenoviral candidates will be included in research and development.

Here, we reported the study of a non-replicating chimpanzee adenovirus-based vaccine against the SARS-CoV-2 Delta variant, which is one of the variants of concern (VOCs). We evaluated the humoral and cellular immune responses elicited by the candidate vaccine in mice. The induction of mucosal immunity and protective efficacy of the candidate vaccine against the Delta variant challenge in mice have also been examined.

## 2. Materials and methods

### 2.1. Cells and viruses

HEK293 cells were purchased from the American Type Culture Collection (ATCC). HEK-293T-ACE2 cells were purchased from YEASEN (Cat. No: 41107ES03, China). Cells were cultured in Dulbecco’s modified Eagle’s medium (Gibco, USA) containing 10% inactivated fetal bovine serum (Gibco, USA) and were cultured at 37 °C and 5% CO_2_. The SARS-COV-2 pseudoviruses were purchased from Beijing Tiantan Pharmacy Biological Research & Development Company. The SARS-COV-2 variants were provided by Wuhan Institute of Virology, Chinese Academy of Sciences (CAS), serial number of the Delta variant is 1633.06.IVCAS 6.7593.

### 2.2 Reagents and kits

The Polyethylenimine Linear (PEI) was purchased from YEASEN (Cat. No: 40816ES02, China) and was used for transfection. The antigen expression was examined by SARS-CoV2 (2019-nCoV) Spike RBD ELISA KIT (Cat. No: KIT40592, SinoBiological, China) following the manufacturer’s instruction. The rescued rAds were purified using Adeno-X Maxi Purification Kit (Cat. No: 631533, Takara, Japan).

### 2.3. Construction of the recombinant viruses

The method to develop a rAd has been described previously (Zhang et al., 2017). In our study, the designed E1/E3 deleted and E4 changed ChAd3 (GenBank ID: CS138463) genome (JS1), about 33 kb, was divided into seven segments according to different restriction sites and was cyclized by the additional pUC57 backbone. It shows as pUC57-*Pac* I-(nt 1-7034)-*Sbf* I-(nt 7043-10666)-*Xba* I-(nt 10673-12532)-*Mlu* I-(nt 12539-19096)-*Cla* I-(nt 19103-25736)-*Asc* I-(nt 25745-33975)-*Eco*R I-(nt 33982-37741)-*Pme* I-pUC57. The seven DNA segments were chemical synthesized by Sangon Biotech and Generay Biotech, and were then ligated head-to-tail with each other sequentially using T4 DNA ligase (NEB, USA). The Delta-*S* sequence was designed according to previous reports (Bos et al., 2020; Liu et al., 2021b; Planas et al., 2021) and was synthesized by Tsingke Biotech. The CMV promoter and BGH poly(A) signal sequences were obtained from pcDNA3.1(+) plasmid by PCR, and were then ligated with the Delta-*S* sequence by overlapping PCR. Meanwhile, I-*Ceu* I and PI-*Sce* I were added to both ends of the CMV-*S*-BGH sequence by PCR. The JS1 plasmid was linearized by a double enzyme digestion with I-*Ceu* I and PI-*Sce* I, and the CMV-*S*-BGH sequence was inserted into JS1 genome by T4 DNA ligase. Finally, HEK293 cells were transfected with JS1 or JS1-delta plasmid using PEI to rescue the recombinant virus.

### 2.4. Animals

Specific pathogen-free (SPF) female BALB/c mice were obtained from Beijing Vital River Laboratory Animal Technology Co., Ltd. SPF H11-K18-hACE2 mice, which is a type of human angiotensin-converting enzyme 2 (hACE2) transgenic C57BL/6J mice, were obtained from Gempharmatech Co., Ltd.

### 2.5. Purification of SARS-CoV-2 Delta-S protein

SARS-CoV-2 Delta-S protein (1-1141aa) with a C-terminal 6× His tag was expressed in HEK293 cells. In brief, adherent HEK293 cells were transfected with recombinant pcDNA3.1-Delta S-His (1-1141) using PEI and incubated in 37 °C, 5% CO_2_ for 72 h. The supernatant was harvested by centrifugation and purified by Ni-NTA affinity chromatography.

### 2.6. Enzyme linked immunosorbent assay (ELISA)

This method has been described previously (Wu et al., 2020). In brief, to measure the titers of SARS-CoV-2-specific IgG or IgA, 96-well clear polystyrene microplates (Corning, USA) were coated with 100 ng/well Delta-S1 protein (GenScript, China) or 250 ng/well purified Delta-S protein in PBS and incubated at 4 °C overnight. After being washed with PBST for 2-4 times, the plates were blocked at 37 °C for 1 h with 5% skim milk in PBST and washed again. Then, the plates were added with 100 μL serially diluted sera or BALFs (bronchoalveolar lavage fluids) or saliva and incubated at 37 °C for 1 h. Following washes, HRP-conjugated Affinipure Goat Anti-Mouse IgG (H+L) (Proteintech, USA, 1:5000 dilution) or Goat Anti-Mouse IgA-HRP (SouthernBiotech, USA, 1:5000 dilution) was added to the plates and incubated for 1 h at 37 °C. The assays were developed with TMB substrate (Sigma, USA), stopped by the addition of 2M H_2_SO_4_ and were measured at 450 nm/630 nm. The endpoint titer was defined as the highest dilution yielding an absorbance >2-fold over the values of negative control.

### 2.7. Neutralization assay

This method has been described previously (Wu et al., 2020). In brief, to test SARS-CoV-2 PNAbs, serial dilutions of heat-inactivated sera were mixed with 1000 TCID_50_ of the Delta variant pseudovirus, incubated for 1 h at 37 °C and added to HEK-293T-ACE2 cells in duplicate in 96-well microplate. The uninfected cells served as a negative control, and the cells infected with pseudovirus-only served as a positive control. Cells were lysed 24 h later and luciferase activity was measured. The half maximal effective concentration (EC_50_) were calculated using Reed Muench method. To test SARS-CoV-2 NAbs, serial dilutions of heat-inactivated sera were incubated with 70 PFU of the indicated SARS-COV-2 variants (isolated and stored by Wuhan Institute of Virology, CAS) at 37 °C for 1 h. The mixtures were added to pre-plated Vero E6 (ATCC) cell monolayers in 96-well microplate and cultured for 3-4 days. NAb titers were calculated by Reed Muench method to estimate the serum dilution required for a 50% cytopathic effect (CPE) reduction.

### 2.8. ELISpot

SARS-CoV-2-specific cellular immune responses were assessed by the IFNγ and IL-2 ELISpot Kit (Cellular Technology Limited, USA) following the manufacturer’s instructions. In brief, 1×10^6^ of mice splenic lymphocytes were stimulated with Delta-S protein in a pre-coated ELISpot plate for 16 h at 37 °C. After being washed twice, the plates were added with Anti-murine IFNγ Detection Solution or Anti-murine IL-2 Detection Solution and incubated at room temperature for 2 h. The plates were then added with Tertiary Solution and Blue Developer Solution. Finally, the spots were counted with ImmunoSpot Analyzer (S6 Universal, Cellular Technology Limited, USA).

### 2.9. Quantitative real time PCR (qPCR)

SARS-COV-2 viral RNA was extracted using TRI Reagent (Sigma, USA). Reverse transcription was performed using HiScript II Q RT SuperMix for qPCR (Vazyme, China). Subsequently, 5 μL cDNA was added into a 25 μL qPCR reaction containing ChamQ SYBR qPCR Master Mix (Vazyme, China). The primers designed to target the nucleocapsid (N) protein of SARS-CoV-2 were (forward) 5’-GGG GAA CTT CTC CTG CTA GAA T-3’ and (reverse) 5’-CAG ACA TTT TGC TCT CAA GCT G-3’. The samples were run in triplicate on an ABI 7900 Real-Time System (Applied Biosystems, Thermo, USA). The following cycling conditions were performed: 1 cycle of 50 °C for 2 min, 1 cycle of 95 °C for 10 min, and 40 cycles of 95 °C for 15 sec and 58 °C for 1 min. Finally, the amount of viral RNA was normalized to the standard curve from a plasmid containing the full-length SARS-CoV-2 *N* gene.

### 2.10. Immunofluorescence (IF) and hematoxylin-eosin staining (H&E)

The lung tissues were immersed in 10% neutral buffered formalin (Sigma, USA) for 24 h. After the formalin fixation, the tissues were placed in 70% ethanol (Merck, Germany) and subsequently embedded with paraffin. Tissue sections (5 μm thick) were prepared for H&E staining or immunofluorescence staining using SARS-CoV/SARS-CoV-2 Nucleocapsid Antibody (SinoBiological, China). Images were collected under a Pannoramic MIDI system (3DHISTECH, Thermo, USA) using Pannoramic scanner software and analyzed by ImageJ (National Institutes of Health, NIH).

### 2.11. Statistical analysis

Data were presented as the mean ± standard error of the mean. Statistics were performed using the Mann-Whitney U test or one-way ANOVA or two-way ANOVA. Tests were performed using GraphPad Prism software (Version 8). P<0.05 was considered statistically significant (* p<0.05, ** p<0.01).

## 3. Results

### 3.1. Construction and identification of JS1-delta

To develop a prophylactic vaccine against the SARS-CoV-2 Delta variant, we constructed a recombinant replication-incompetent adenovirus carrying optimized Delta spike protein (S)-coding sequence, named JS1-delta. Chimpanzee adenovirus type 3 (ChAd3, GenBank ID: CS138463) was chosen to delete E1 region (nt 589-3544) and E3 region (nt 28310-32653). Meanwhile, ChAd3 E4 region (nt 34698-37314) was replaced by the E4 region of hAd5 (GenBank ID: AY339865, nt 32909-35522) to ensure excellent yield of the rAd on producer cells (Abbink et al., 2007; Havenga et al., 2006; Täuber and Dobner, 2001). Two restriction enzymes, I-*Ceu* I and PI-*Sce* I, were inserted into the original ChAd3 E1 region. This recombinant ChAd3 vector was named JS1 (Fig. 1A). The S-coding sequence from the Delta variant was optimized by altering the codon usages to increase its antigen expression in mammalian cells (Liu et al., 2021b; Planas et al., 2021). At the same time, the furin cleavage site mutations (R682S, R685G) and proline substitutions (K986P, V987P) were introduced in the sequence to stabilize the S protein and prevent syncytium formation (Bos et al., 2020). Then, with the addition of CMV promoter and BGH poly(A) signal, the CMV-*S*-BGH segment was inserted into JS1 between I-*Ceu* I and PI-*Sce* I (Fig. 1B). As shown in Fig. 1C, JS1-delta was successfully rescued and multiplied in HEK293 cells, resulting in efficient expression of S protein. At the same time, S expressions in uninfected cells and cells infected with JS1 were undetectable.

**Fig. 1.**
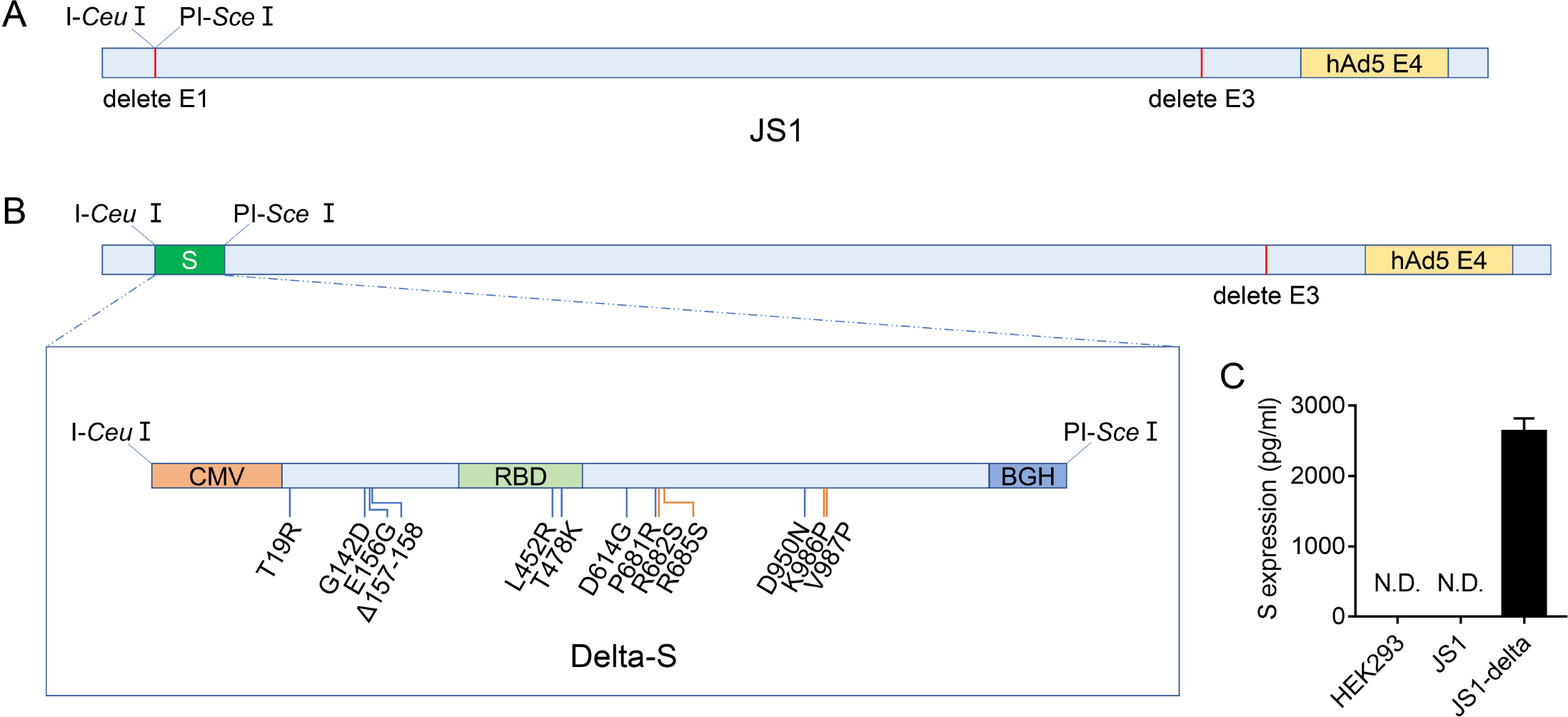
Construction and identification of JS1-delta. **A-B** Schematic of JS1 and JS1-delta genome. S: SARS-CoV-2 spike protein. CMV: cytomegalovirus promoter. RBD: receptor-binding domain of SARS-CoV-2 spike protein. BGH: bovine growth hormone poly(A) signal. hAd5: human adenovirus type 5. The natural mutant sites of Delta S protein were shown as blue lines, and the artificially modified sites were shown as orange lines. **C** HEK293 cells were infected with JS1 or JS1-delta and cultured for 72 hours. Along with the supernatants, cells were frozen and thawed for three times. The cell lysates were collected to detect the expression of S protein by ELISA. N.D. : not detected.

### 3.2. JS1-delta induces potent humoral and cellular immune responses in mice

To assess the immunogenicity of JS1-delta, six-week-old female BALB/c mice (n=5 per group) received a single immunization of 5×10^7^ virus particles (VP) (low dose), 5×10^8^ VP (middle dose), 5×10^9^ VP (high dose) of JS1-delta or 5×10^9^ VP of the control vector JS1 via the intranasal (IN) route at day 0. The titers of Delta-S1-specific IgG, neutralizing antibody (NAb) and cellular immune responses were detected in each group. The serum IgG titers were detectable at week 2 and stayed at a high level until week 8 (Fig. 2A). Similar to the IgG titers, the Delta variant pseudovirus NAb (PNAb) titers were detectable at week 2 and stayed steady from week 4 to week 8 (Fig. 2B). Moreover, all three doses of JS1-delta could elicit potent interleukin-2 (IL-2) response in splenic lymphocytes at week 8, but only the high-dose group induced significant interferon-γ (IFNγ) response (Fig. 2C).

**Fig. 2.**
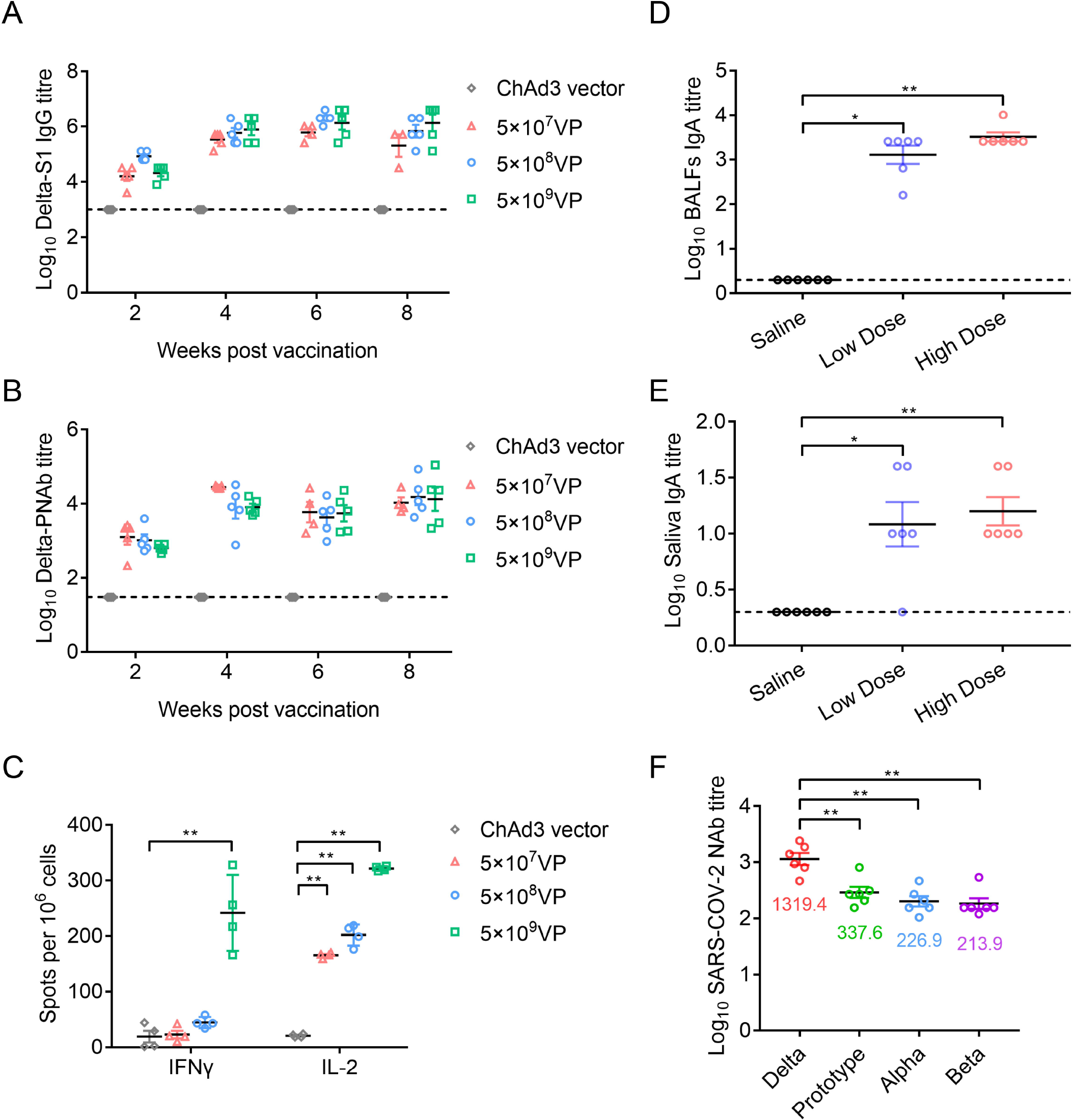
The immunogenicity of JS1-delta in mice. 6-week-old BALB/c mice (n=5 per group) were vaccinated intranasally with the indicated dosages of JS1-delta or the control ChAd3 vector (JS1) at day 0. **A** The titers of S1-specific IgG were assessed at weeks 2, 4, 6 and 8 following vaccination by ELISA. The initial serum dilution is 1:1000. **B** The titers of Delta PNAb were examined by luciferase assay. The initial serum dilution is 1:30. **C** The levels of cellular immune responses at week 8 were detected by ELISpot. 33-day-old H11-K18-hACE2 mice (n=6 per group) were immunized intranasally with 5×10^8^ VP (low dose) and 5×10^9^ VP (high dose) of JS1-delta or the same volume of saline at day 0. At week 4, the S-specific IgA titers in BALFs (**D**) or saliva (**E**) were examined by ELISA. The initial dilution is 1:2. **F** The NAb titers of the high dose group against the indicated SARS-COV-2 variants were detected by CPE calculation.

### 3.3. JS1-delta induces effective mucosal immune responses in hACE2 transgenic mice

To further study the ability of JS1-delta for mucosal immunity induction, the 33-day-old female H11-K18-hACE2 mice (n=6 per group) were immunized intranasally with 5×10^8^ VP (low dose) and 5×10^9^ VP (high dose) of JS1-delta or the same volume of saline as a control at day 0. The Delta-S-specific IgA titers were examined in each group at week 4 post vaccination. As shown in Fig. 2D and E, both low- and high-dose groups elicited efficient S-specific IgA expressions in the bronchoalveolar lavage fluids (BALFs) and the saliva of mice. Meanwhile, the cross-protection of JS1-delta against SARS-CoV-2 prototype, Alpha (B.1.1.7), Beta (B.1.351) and Delta (B.1.617.2) variants was also evaluated. As shown in Fig. 2F, the high-dose group showed high NAb titers against all four SARS-CoV-2 variants at week 4. But compared with those against the Delta variant, the NAb titers against other three variants decreased by about 4-6 folds.

### 3.4. JS1-delta protects mice from the Delta variant infection

To determine the protective efficacy of JS1-delta through the mucosal vaccination, 33-day-old female H11-K18-hACE2 mice were immunized with 5×10^8^ VP (low dose) and 5×10^9^ VP (high dose) of JS1-delta or the same volume of saline by the IN route at day 0. Then, mice were challenged with 1×10^3^ PFU of the Delta variant by nasal drop at day 28 (Fig. 3A). As shown in Fig. 3B and C, all vaccinated mice showed high levels of serum IgG and NAbs against the Delta variant at day 26, which were undetectable in control animals. Whereafter, we monitored the weight change and the survival rate in each group. The evaluating indicators kept normal in all groups until day 3 post-infection (3 dpi), but the infected mice without immunization (saline group) lost weight obviously from 4 dpi and all died at 5 dpi, while other groups remained stable and stayed alive (Fig. 3D and E). Subsequently, mice were euthanized at 6 dpi to detect the viral load in lung tissue and for immunofluorescence or pathological examination. As expected, only the saline group exhibited a significantly high level of viral RNA copies, displayed lots of SARS-CoV-2 N protein positive cells and showed severe histopathological changes in lung tissue (Fig. 3F and G). Taken together, all these results indicated that both low- and high-dose JS1-delta can confer sufficient protection against the Delta variant infection in hACE2 transgenic mice.

**Fig. 3.**
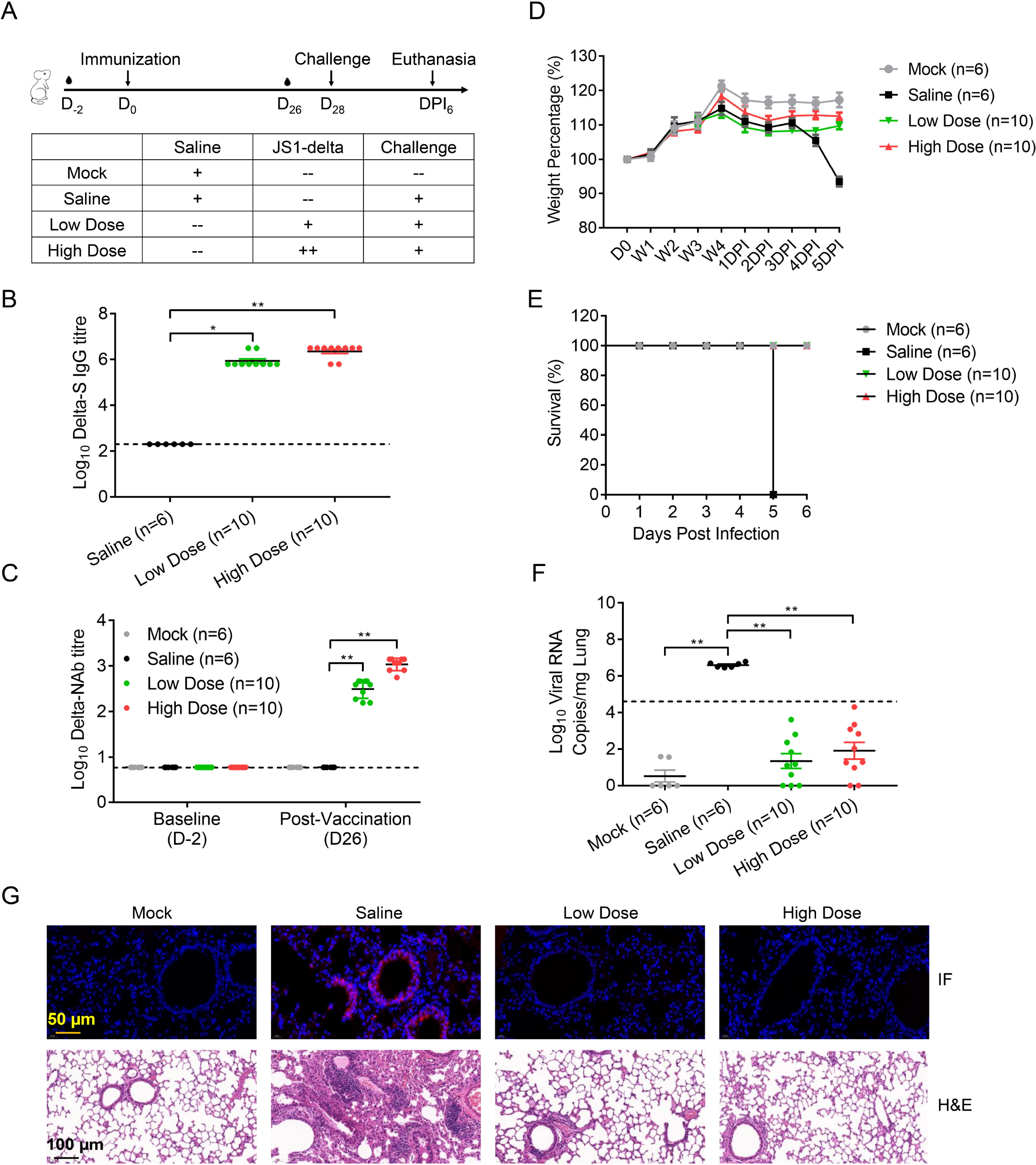
JS1-delta protects mice from the Delta variant infection. **A** Schematic of experimental strategy and grouping. 33-day-old H11-K18-hACE2 mice were immunized intranasally with 5×10^8^ VP (low dose, n=10) and 5×10^9^ VP (high dose, n=10) of JS1-delta or the same volume of saline (n=6) at day 0. Then, the mice were challenged with 1×10^3^ PFU of the Delta variant in a volume of 50 μL by nasal drop at day 28. Serums were collected at day −2 and day 26. **B** The titers of S-specific IgG at day 26 were detected by ELISA. The initial serum dilution is 1:200. **C** The titers of the Delta NAbs were detected by CPE calculation. The initial serum dilution is 1:6. **D** The weight change was monitored every week after vaccination and every day after infection. **E** The survival rate was monitored every day after infection. The mice in saline group died at 5 dpi, the others were sacrificed at 6 dpi. **F** The viral load in lung tissue was detected by qPCR. The lower limit of qPCR detection was set to a Ct (cycle threshold) value equal to 33. **G** SARS-COV-2 N protein expression and histopathological examinations in lungs were examined by immunofluorescence (IF) and hematoxylin-eosin staining (H&E).

## 4. Discussion

Multiple reports have found that the SARS-CoV-2 variants can reduce the neutralization capability of monoclonal antibodies (mAbs) and convalescent plasma, and can resist antibody-mediated immunity elicited by vaccines (Cao et al., 2021a; Cao et al., 2021b; Cele et al., 2021; Li et al., 2021; Liu et al., 2021a; Lustig et al., 2021; Planas et al., 2021; Shen et al., 2021; Tian et al., 2021). As stated by the WHO Technical Advisory Group on COVID-19 Vaccine Composition (TAG-CO-VAC), considering supply and demand of the available vaccines, along with evolution of the virus, repeated booster doses of the original vaccine composition is unlikely to be appropriate or sustainable (WHO, 2022b). Based on the reality that the new variants emerge constantly, it is not enough for vaccines to just reduce hospitalization and mortality, but should be more effective in preventing infection and transmission. In our study, JS1-delta has been demonstrated to protect mice from the Delta variant infection sufficiently (Fig. 3). However, its cross-protection against other variants is attenuated (Fig. 2F), which is consistent with another COVID-19 candidate vaccine JS1-beta (Fig. S1E). These results and reports suggest that a monovalent vaccine used independently may be difficult to handle the complex epidemic situation. Moreover, it is reported that the heterologous prime-boost strategy with COVID-19 vaccines from diverse platforms could be more effective than using a homologous booster (Liu et al., 2021c; Zhang et al., 2021). Therefore, using heterologous boosters or developing multivalent vaccines containing various SARS-CoV-2 variants compositions may be more efficient and practical to fight against the virus.

Replication deficient recombinant adenoviral vectors possess great immunogenicity and can elicit potent cellular immune responses in preclinical and clinical researches, in which hAd5 is the most comprehensively studied (Abbink et al., 2007; Bassett et al., 2011; Catanzaro et al., 2006; Quinn et al., 2013; Sullivan et al., 2003). However, hAd5 shows high seroprevalence in global populations due to natural infection, especially in the developing world, and high level of the pre-existing immunity could result in decreased efficacy of the hAd5-based vaccines (Abbink et al., 2007; Quinn et al., 2013). Meanwhile, considering that HEK293 remains a major producer cell line for adenovirus development and manufacturing, replication-competent adenovirus (RCA) contamination is a significant trouble for large-scale production and clinical application of the hAd5-derived vectors (Kovesdi and Hedley, 2010; Lusky, 2005). To circumvent these limitations, considerable efforts have been made to develop new adenoviral vectors from lower seroprevalence hAds (such as hAd26, hAd28 and hAd35) (Abbink et al., 2007; Kahl et al., 2010; Vogels et al., 2003) or from non-human primate sources (such as ChAdY25, Simian Ad36, ChAd63 and ChAd68) (Dicks et al., 2012; Hassan et al., 2020; O’Hara et al., 2012; Wang et al., 2019). ChAd3 is one of the most promising candidates and shows well safety and tolerance in clinical trials targeting hepatitis C virus and Ebola virus (Barnes et al., 2012; De Santis et al., 2016; Kelly et al., 2016; Ledgerwood et al., 2017). In our study, we demonstrated for the first time that a ChAd3-derived COVID-19 vaccine can induce potent humoral and cellular immune responses, protect mice from the Delta variant challenge effectively. At the same time, another vaccine candidate JS1-beta targeting the Beta (B.1.351) variant shows favorable immunogenicity in mice (Fig. S1), and JS1-omicron targeting the Omicron (B.1.1.529) variant is also under testing. The longevity of antigen-specific immune responses and the influence of anti-vector immunity induced by the vaccines, along with the safety evaluation and the cooperation with different boosters, will be further investigated. What calls for special attention is that some vaccines can cause rare adverse events. It is reported that the AstraZeneca and Janssen vaccines are related with thrombosis with thrombocytopenia syndrome (TTS), immune thrombocytopenic purpura (ITP) and Guillain-Barre syndrome (GBS) (CDC, 2022a; Cines and Bussel, 2021; McGonagle et al., 2021; Rahman et al., 2022; WHO, 2022c). Meanwhile, mRNA vaccines, including BNT162b2 developed by Pfizer and mRNA-1273 developed by Moderna, can cause myocarditis, pericarditis and anaphylaxis (CDC, 2022a; Rahman et al., 2022; WHO, 2022c). The pathogenesis of these rare adverse events is not yet clear, but as stated by the Centers for Disease Control and Prevention (CDC), the benefits of COVID-19 vaccination continue to outweigh any potential risks and the safety of COVID-19 vaccines will be under continuous monitoring (CDC, 2022b).

Recently, mucosal vaccines against SARS-CoV-2 have raised increasing interests due to their important advantage in preventing virus replication in the upper respiratory tracts, which is essential to interrupt pulmonary infection and person to person transmission (Feng et al., 2020; Hassan et al., 2020; Lavelle and Ward, 2021; Wu et al., 2020). Compared with intramuscular injection, the IN vaccination can elicit not only comparable systemic immunity but also additional pathogen-specific mucosal immunity, resulting in a first line of defense against the virus invasion (Feng et al., 2020; Hassan et al., 2020; Wu et al., 2020). In our study, IN vaccination of JS1-delta induces S-specific IgA expressions efficiently in both BALFs and saliva of the hACE2 transgenic mice (Fig. 2D and E). Using SARS-CoV-2 as a model, our findings both *in vitro* and *in vivo* highlight that the non-replicating ChAd3-derived vector administrated in the IN route represents a powerful platform for rapid vaccine development against respiratory virus infection.

## 5. Conclusions

In summary, we developed and tested a recombinant ChAd3-derived viral vaccine against the SARS-COV-2 Delta variant infection. With further preclinical and clinical researches, this kind of vaccine is hopeful to provide a new option to fight against the COVID-19 pandemic.

## Data availability

All data and materials are available in the main text or the supplementary materials.

## Ethics statement

The experiments involving animals were approved by and carried out in accordance with the guidelines of the Institutional Experimental Animal Welfare and Ethics Committee.

## Author contributions

Yifei Yuan: data curation, formal analysis, writing-original draft. Xing Gao: project administration, resources. Fengfeng Ni: data curation, methodology. Wenbo Xie: data curation, methodology. Wenbin Fu: data curation, methodology. Gaoxia Zhang: resources, supervision. Huimin Hu: data curation, methodology. Yuncheng Li: data curation, methodology. Qinxue Hu: formal analysis, resources, supervision. Chuanmengyuan Huang: methodology. Bo Liu: methodology, supervision. Yalan Liu: formal analysis, writing-review & editing. Qiong Shen: methodology, writing-review & editing. Min Liang: conceptualization, writing-review & editing.

## Acknowledgments

We are particularly grateful to Prof. Bing Sun and Dr. Xiaoyu Sun (Center for Excellence in Molecular Cell Science, CAS) for the technical assistance in IgG ELISA and PNAb assay. This research did not receive any specific grant from funding agencies in the public or not-for-profit sectors.

