## Supplementary material for "A novel intranasal administration adenoviral vector-based platform for rapid COVID-19 vaccine development": Fig. S1

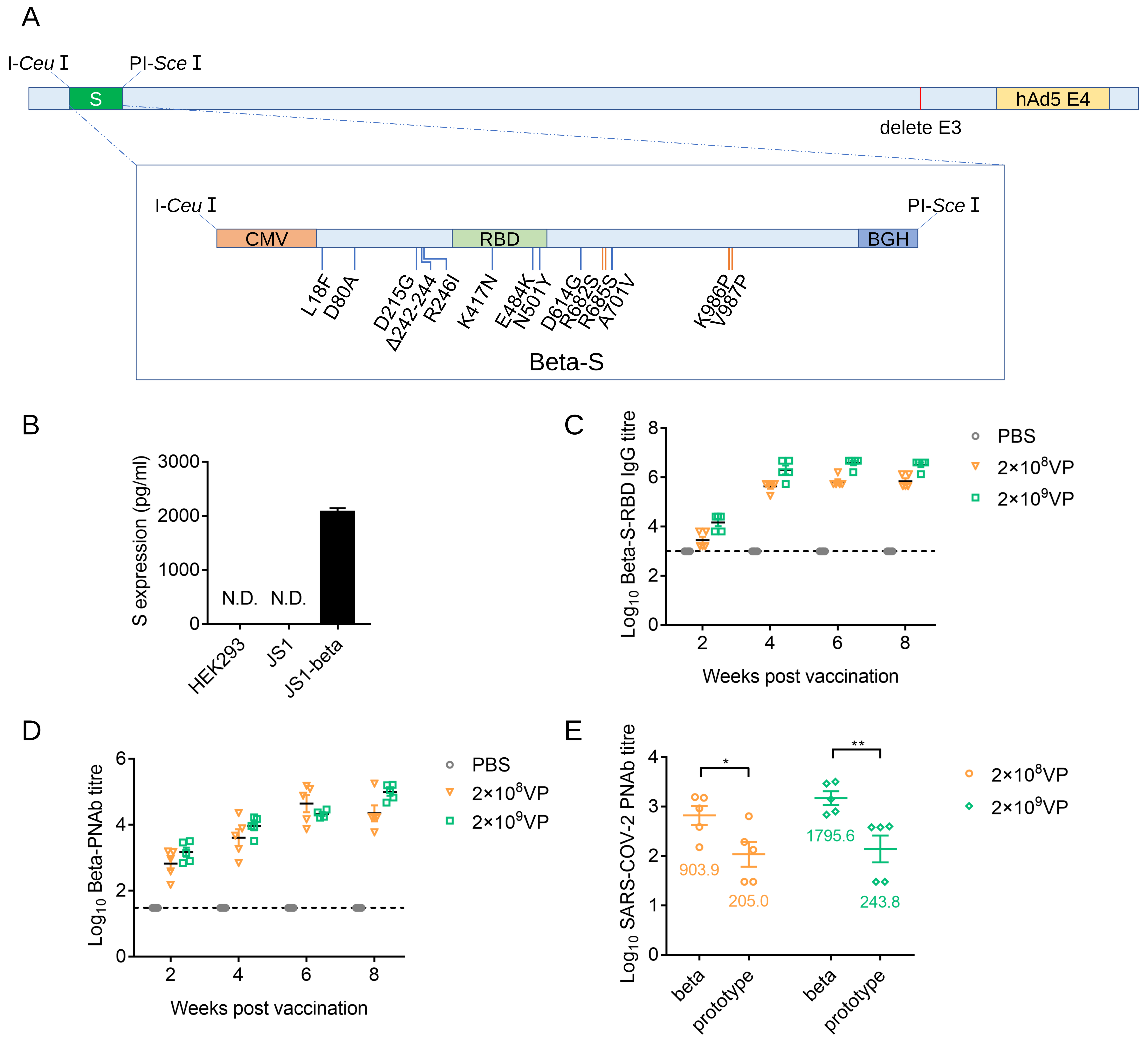


**Fig. S1.** The immunogenicity of JS1-beta in mice. **A** Schematic of JS1-beta genome. S: SARS-CoV-2 spike protein. CMV: cytomegalovirus promoter. RBD: receptor-binding domain of SARS-CoV-2 spike protein. BGH: bovine growth hormone poly(A) signal. hAd5: human adenovirus type 5. The natural mutant sites of Beta S protein were shown as blue lines, and the artificially modified sites were shown as orange lines. **B** HEK293 cells were infected with JS1 or JS1-beta and cultured for 72 hours. Along with the supernatants, cells were frozen and thawed for three times. The cell lysates were collected to detect the expression of S protein by ELISA. N.D. : not detected. **C** 6-week-old BALB/c mice (n=5 per group) were vaccinated intranasally with JS1-beta or the same volume of PBS using the indicated dosages at day 0. The titers of Beta-RBD-specific IgG were assessed at weeks 2, 4, 6 and 8 by ELISA. The initial serum dilution is 1:1000. **D** The titers of Beta PNAb were examined by luciferase assay. The initial serum dilution is 1:30. **E** The cross-protection against the indicated SARS-COV-2 variants at week 2 was detected by luciferase assay.
